# Zika Virus in the Human Placenta and Developing Brain: Cell Tropism and Drug Inhibition

**DOI:** 10.1101/058883

**Authors:** Hanna Retallack, Elizabeth Di Lullo, Carolina Arias, Kristeene A. Knopp, Carmen Sandoval-Espinosa, Matthew T. Laurie, Yan Zhou, Matthew Gormley, Walter R. Mancia Leon, Robert Krencik, Erik M. Ullian, Julien Spatazza, Alex A. Pollen, Katherine Ona, Tomasz J. Nowakowski, Joseph L. DeRisi, Susan J. Fisher, Arnold R. Kriegstein

**Affiliations:** Department of Biochemistry and Biophysics, University of California San Francisco, San Francisco, CA 94158, USA; Eli and Edythe Broad Center of Regeneration Medicine and Stem Cell Research, University of California, San Francisco, San Francisco, CA 94143, USA; Department of Neurology, University of California, San Francisco, San Francisco, CA 94158, USA; Center for Reproductive Sciences, University of California San Francisco, San Francisco, CA 94143, USA; Department of Obstetrics, Gynecology, and Reproductive Sciences, University of California San Francisco, San Francisco, CA 94143, USA; Department of Anatomy, University of California San Francisco, San Francisco, CA 94143, USA; Department of Ophthalmology, University of California San Francisco, San Francisco, CA 94122, USA; Department of Neurological Surgery, University of California-San Francisco, San Francisco, California 94143, USA; Howard Hughes Medical Institute, Chevy Chase, MD, USA.

## Abstract

The rapid spread of Zika virus (ZIKV) and its association with abnormal brain development constitute a global health emergency. Congenital ZIKV infection produces a range of mild to severe pathologies, including placental damage and microcephaly. However, the placenta’s role in viral transmission and the mechanisms of microcephaly have not been addressed in primary human tissues. Moreover, there is an urgent need for drugs that can prevent developmental defects following infection. Here, we identify the placental and brain cell populations most susceptible to ZIKV infection, provide evidence for a mechanism of viral entry, and show that a commonly used antibiotic protects cultured brain cells by inhibiting viral proliferation. In the early gestation placenta, the virus readily infected trophoblast subpopulations that are in direct contact with maternal blood and uterine cells, suggesting routes of ZIKV transmission to the embryo and fetus. In the brain, ZIKV preferentially infected neural stem cells, astrocytes, and microglia, whereas neurons were less susceptible to infection. These findings suggest mechanisms for microcephaly and other pathologic features of infants with congenital ZIKV infection that are not explained by neural stem cell infection alone, such as calcifications in the cortical plate and brain abnormalities caused by third trimester infection. Blocking a putative viral entry receptor, AXL, which is highly enriched in the infected placenta and brain cell types, reduced ZIKV infection of astrocytes *in vitro*. In a glial cell line, the macrolide antibiotic, azithromycin, inhibited viral proliferation and viral-induced cytopathic effects at clinically relevant concentrations. Our characterization of infection in primary human tissues clarifies the pathogenesis of congenital ZIKV infection and provides critical context for interpreting results from model systems. Further work on azithromycin and related compounds may yield additional therapeutic strategies to safely alleviate or prevent the most severe consequences of the epidemic.

---

Understanding the cell types initially vulnerable to ZIKV infection may help to reveal routes of viral spread, mechanisms underlying fetal abnormalities, and relevant cellular targets for testing therapeutic compounds. The enriched expression of candidate flavivirus entry proteins in specific cell populations suggests that reduced cellular models may not capture the diversity of cell types that could be vulnerable to infection. To study ZIKV infection in the context of the complexities of placental and brain architecture *in vivo,* we exposed organotypic tissue cultures of the placenta and cerebral cortex from relevant developmental stages to three strains of ZIKV: Cambodia 2010 FSS13025 (ZIKV-CAM), Brazil 2015 SPH2015 (ZIKV-BR), and Puerto Rico 2015 PRVABC59 (ZIKV-PR), see Methods.

In clinical reports, viral infection of the placenta and resulting pathology indicate that some fetal abnormalities, including growth restriction, may be attributable to infection of the placenta itself rather than to transplacental passage of the virus and infection of the embryo and fetus^1^. However, recent analyses suggest that trophoblasts isolated from placentas at term are poorly permissive to ZIKV infection^2,3^. To explore viral transmission during early pregnancy when the formative stages of brain development take place, we utilized an organotypic culture system of chorionic villus explants from first and second trimester human placentas. These organoids contain the trophoblast (TB) cell types that are in direct contact with maternal blood and/or cells of the uterus: multi-nucleated syncytiotrophoblasts (STBs) that form the villus surface and cell columns, which contain mononuclear cytotrophoblasts (CTBs) that are destined to invade the uterus and its resident blood vessels (diagrammed in **Extended Data** Fig. 1). The explants were cultured for 24 h prior to infection, which enabled re-establishment of their basic architecture, including cell column outgrowths.

Immunostaining with an antibody against the flavivirus envelope protein enabled visualization of infection. STBs and CTBs were distinguished by cytokeratin (CK) expression. Twenty-four hours post-infection (hpi) with ZIKV-BR we observed patches of STBs at the villus surface that were immuno-positive for the viral protein. Occasionally the underlying CTBs also reacted with the antibody (Fig. 1a). The explant model includes CTB outgrowths, the *in vitro* equivalent of cell column extensions, that were also infected (Fig. 1b). In some areas, there was evidence of infected CK-negative stromal elements (Fig. 1b, upper left). During the first trimester, the same pattern of ZIKV infection was observed with all the strains that were tested (**Extended Data** Fig. 2). Freshly isolated CTBs purified from first or second trimester placentas were also susceptible to infection (**Extended Data** Fig. 3). Second trimester explants had many fewer foci of infection than the first trimester cultures, but the pattern of STB and CTB staining with anti-flavivirus envelope protein was the same (Fig. 1c). At this stage, cell columns, which are largely depleted of CTBs, no longer form in culture and thus could not be analyzed. The patterns of gestational-age related changes in susceptibility to infection mirrored the decline in expression of the putative ZIKV receptor, AXL, in CTBs and STBs from first to second trimester (**Extended Data** Fig. 4 and 5). Interestingly, infection of STBs and CTBs was associated with disorganized cytokeratin, revealing possible evidence of cellular damage (Fig. 1a). Furthermore, immunoblotting showed that infection was accompanied by increased expression of AXL and markers of autophagy, LC3B and p62 (**Extended Data** Fig. 3). Together, these data showed that the TB cell types that are in direct contact with maternal blood and the uterine wall are susceptible to ZIKV infection during the first half of pregnancy, indicating vulnerability at stages relevant to fetal abnormalities and suggesting routes of placental infection across floating villi or via the cell columns of anchoring villi that are attached to the uterine wall.

**Figure 1.**
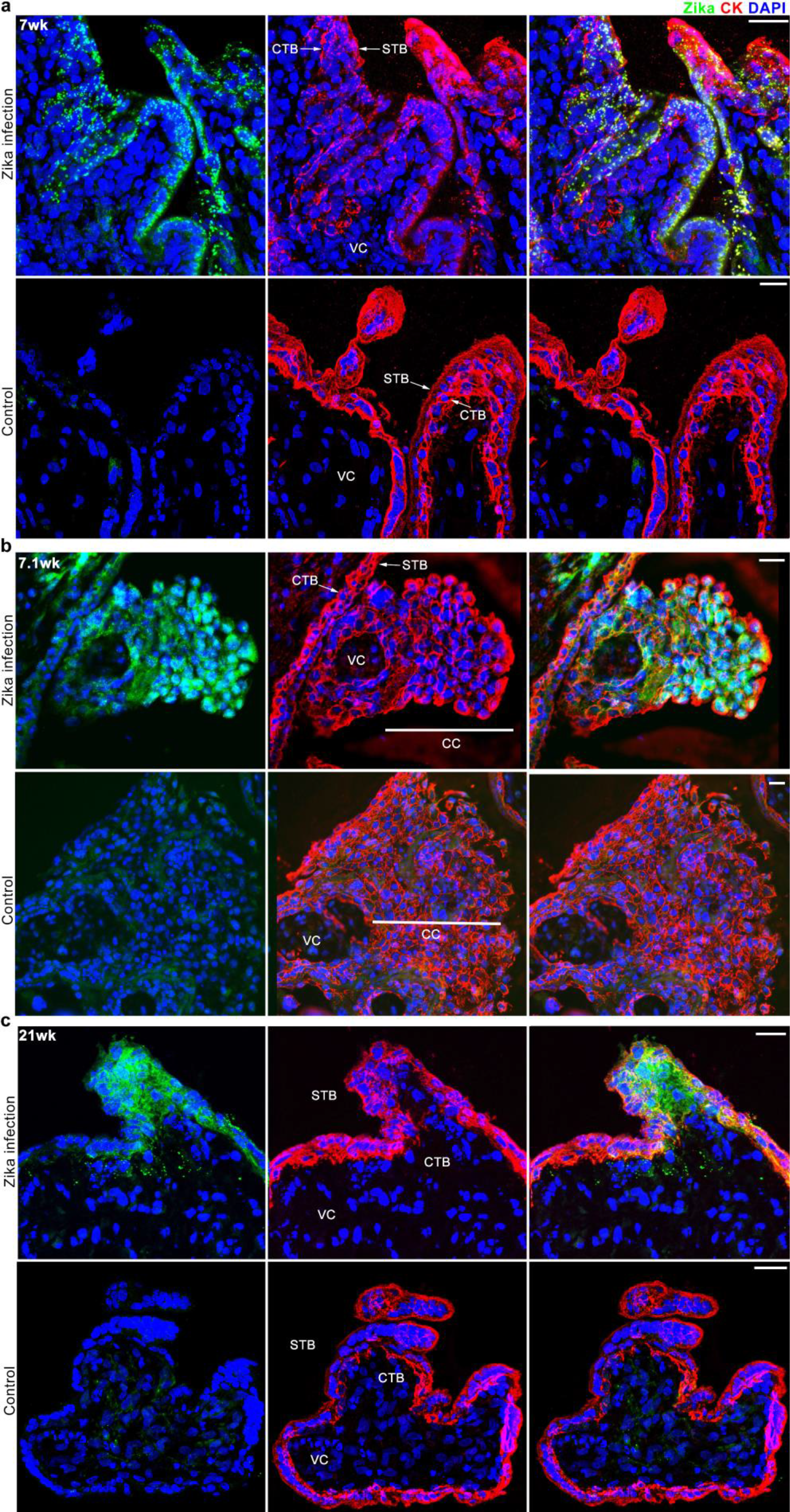
Tropism of ZIKV in human chorionic villus explants during the first and second trimesters of pregnancy. The explants were cultured for 24 h prior to infection with ZIKV-BR. Cytokeratin (CK) immunoreactivity (red) confirmed syncytiotrophoblast (STB) and cytotrophoblast (CTB) identity. The nuclei were stained with DAPI. **a**, (7 wks) At 24 hpi, patches of STBs at the villus surface reacted with anti-flavivirus envelope protein (green). In some locations, the underlying CTBs were immuno-positive. **b**, (7.1 wks) There was also evidence of CTB cell column (CC) infection. CK-negative stromal elements also exhibited immunoreactivity. **c**, (21 wks) As pregnancy advanced, the same pattern of infection was observed at the villus surface, but overall there were fewer foci (data not shown). The control cultures were mock infected. Images were acquired by confocal **(a, c)** and fluorescence (**b**) microscopy. Scale bars, 25 µm.

After crossing the placenta, ZIKV is neurotropic, as established by the detection of viral RNA and flavivirus particles in post-mortem brain tissue of infants with microcephaly^4^. Recent studies have begun to study the consequences of infection in permissive mouse strains and in neural cells dissociated from primary tissue^5^ or derived *in vitro* from human induced pluripotent stem (iPS) cells^6–9^. In the developing mouse brain, ZIKV was found in radial glia, the neural stem cells of the developing brain, and in neurons^10^. ZIKV infection of human iPS-derived neural cells and cerebral organoids highlighted the selective vulnerability of progenitor cells resulting in their apoptotic cell death^6,9^. A study of isolated primary human neural progenitor cells showed ZIKV was initially cytopathic, but that viral replication persisted for weeks in surviving cells^5^. However, none of these models fully recapitulates the developmental events and cell types present during human brain development, and it is unclear how well the models represent ZIKV-induced pathology in human fetuses. Previous studies suggested that enriched expression of AXL by specific cell types could confer vulnerability of those cells to ZIKV entry^11^. Based on the expression of the candidate entry factor AXL, we predicted that radial glia, astrocytes, microglia, and cells lining blood vessels may be particularly vulnerable to infection^12^. Therefore, we next investigated the infectivity of ZIKV in developing human brain using organotypic cultures from primary tissue that preserve tissue architecture, cell behaviors, and many aspects of cell diversity.

In tissue samples from peak stages of neurogenesis (13-16 post-conception week (pcw)), we observed high levels of infection in the ventricular and subventricular zones where radial glia and newborn neurons reside (Fig. 2a). We found that the virus preferentially infected radial glia cells across the germinal zone (Fig. 2b). A minor fraction of cells positive for ZIKV at these stages included postmitotic neurons, that may represent neurons generated from infected radial glia. Interestingly, we observed clusters of infected radial glia in the germinal zone (Fig. 2b), which may reflect spread of the virus from initially infected cells to neighboring radial glia. We observed similar patterns of infection across strains (**Extended Data** Fig. 6), but we did not observe appreciable cell death (**Extended Data** Fig. 7). The finding of preferential radial glia infection and survival in organotypic slices is consistent with experiments in cultured primary human neural stem cells^5^ and developing mouse cortex^10^. Recent studies of dissociated and *in vitro* derived neural stem cells also reported cell death in neural stem cells^6,9,13^not observed here. These observations could reflect differences in maturation stage and culture conditions of cells or in the viral titer and duration of exposure. Even without immediately causing cell death, ZIKV infection of radial glia cells could affect cell cycle progression^10,13^, neuronal differentiation, or the migration and survival of newborn daughter neurons. These mechanisms would be comparable to many genetic causes of microcephaly and lissencephaly that affect radial glia progenitor cells and neuronal migration^14^.

**Figure 2.**
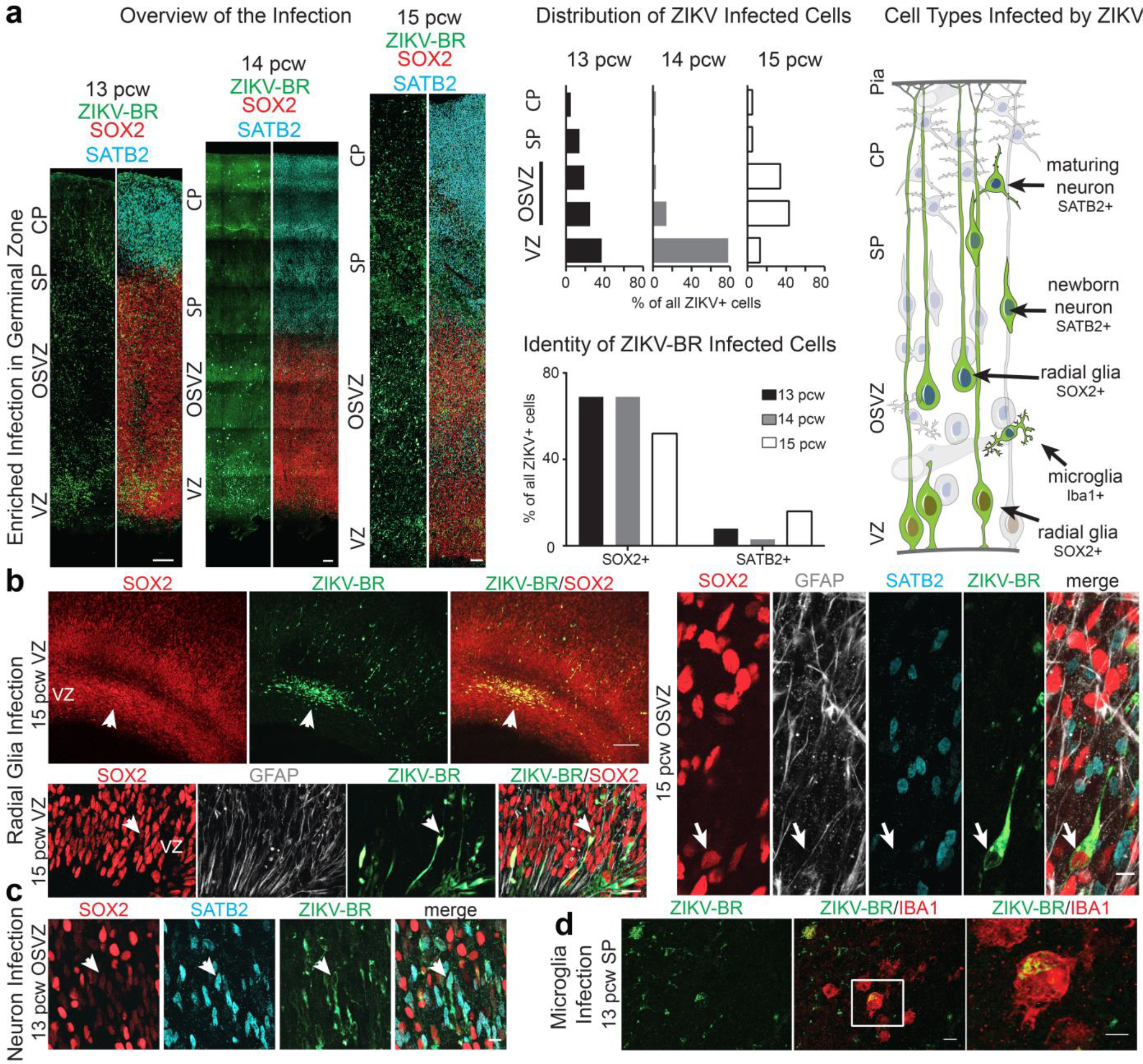
Tropism of ZIKV in the developing human brain at mid-neurogenesis. **a,** Organotypic human brain slices (*n* = 3 independent biological replicates) were exposed to ZIKV-BR and cultured for 72 h. Cell types infected with ZIKV were identified by immunohistochemistry. Immunostaining for ZIKV envelope protein revealed enriched infection in germinal zones, ventricular zone (VZ) and outer subventricular zone (OSVZ). Images show low magnification overview of the infected slices. Scale bar 100 µm. SP ‒ subplate, CP ‒ cortical plate, pcw ‒ post‑conception weeks. Top bar charts represent spatial distribution of ZIKV infected cells. Bottom bar chart represents quantification of the fraction of infected cells that expressed thxse neuronal marker SATB2 or radial glial marker SOX2. Schematic summarizes cell types infected by ZIKV (green) in the developing human brain around mid-neurogenesis. **b**, Radial glia, the neural stem cells of the developing brain, are vulnerable to ZIKV infection. Upper left panels show low magnification view of a representative pattern of ZIKV infection in the cortical VZ. Note a high density cluster of infected cells near the VZ (arrow). Scale bar 100 µm. Bottom left panel shows high magnification view of the cortical VZ. Arrow indicates a single radial glia cell infected with ZIKV. Scale bar 10 µm. Right panels represent high magnification view of the OSVZ with arrow indicating an OSVZ radial glia cell infected with ZIKV. Scale bar 10 µm. **c**, Example of a neuron infected with ZIKV (arrow). Scale bar 20 µm. **d**, ZIKV infects microglia, the immune cells of the brain. d, Panel shows ZIKV infected microglia with amoeboid morphology typical of activated microglia. Scale bar 10 µm.

At later stages of development (after 18 pcw), we observed broad infection of radial glia in the germinal zones. In addition, we observed high infectivity in the cortical plate and subplate where the two principal descendants of radial glia, neurons and astrocytes, migrate during the generation of the cerebral cortex (Fig. 3). Among cortical plate cells, we observed a high rate of infection of astrocytes (Fig. 3a,d,e), consistent with the high level of AXL expression in this cell type^12^. We also observed infection of oligodendrocyte precursor cells (OPCs) and microglia, but there was limited infection of neurons or evidence of exacerbated cell death across developmental stages at 72 hpi (Fig. 3b,c). No major differences in cell type-specific infectivity were observed across the three strains of ZIKV in our short term cultures (**Extended Data** Fig. 7). The high levels of infection in astrocytes, many of which contact microcapillaries, could link our understanding of initial infection with clinical findings of cortical plate damage. For example, after prolonged infection, viral production in astrocytes could lead to a higher viral load in the cortical plate causing infection of additional cortical cell types, and astrocyte loss could lead to inflammation and further damage. Widespread cell death of cortical plate populations is expected based on clinical reports of band-like calcifications in the cortical plate, cortical thinning, and hydrocephalus^4,15^.

**Figure 3.**
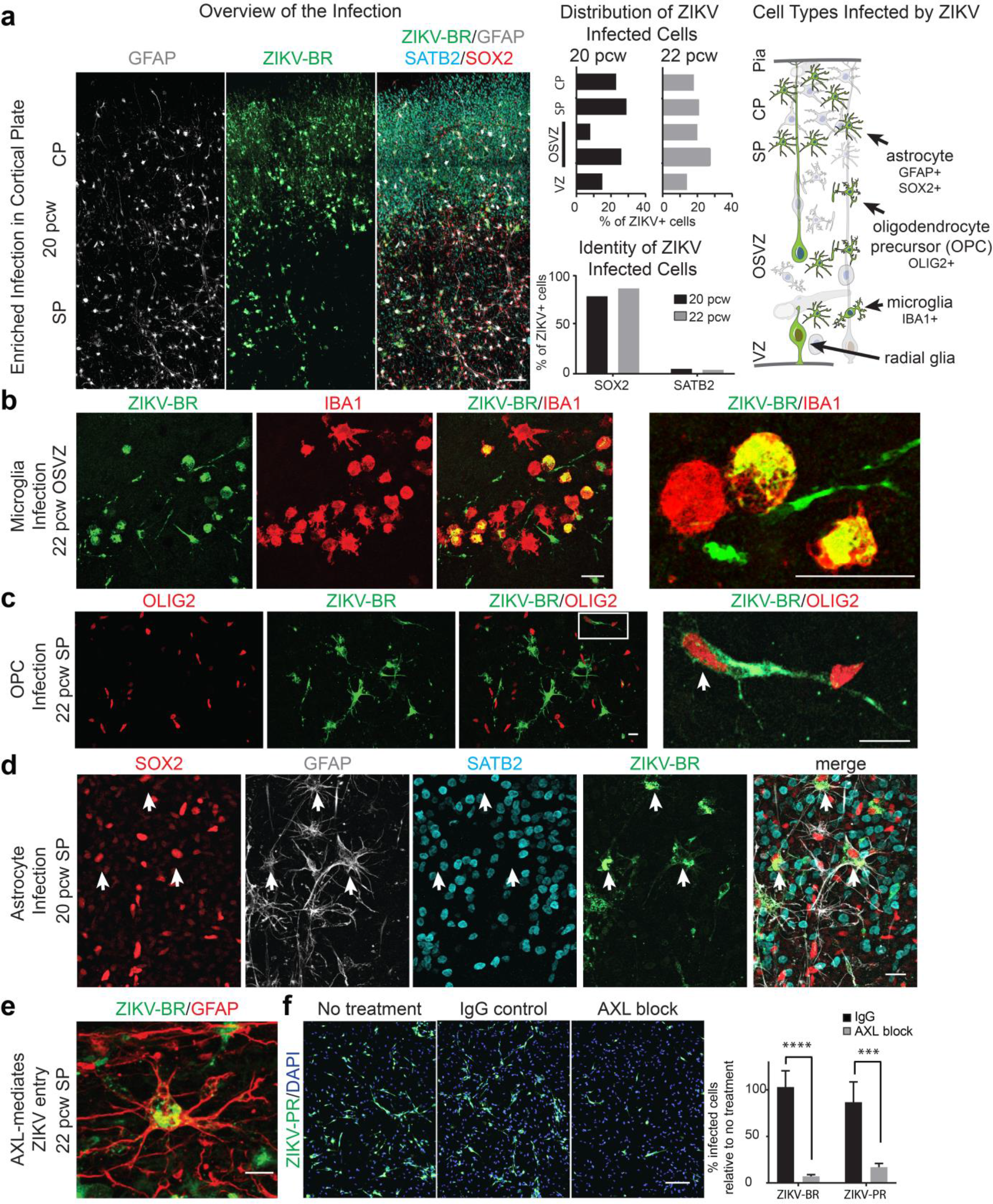
Astrocytes are particularly vulnerable to ZIKV infection. **a**, Image shows low magnification overview of a representative pattern ZIKV infection in human cortical plate at stages corresponding to early astrogliogenesis. Scale bar 100 µm. ZIKV infectivity was analyzed using immunohistochemistry (see Methods). Top bar chart illustrates spatial distribution of infected cells. Bottom bar chart shows the percentage of ZIKV infected neuronal (SATB2+) and glial cells (SOX2+). Schematic summary of ZIKV infected cells (green). **b**, ZIKV infects microglia in late second trimester brain. Left hand panels show low magnification overview of immunostaining of infected microglia (red). Right, high magnification showing infected microglia. Scale bar 50 µm. **c**, Oligodendrocyte Precursor Cells (OPCs) are infected by ZIKV. Left panels show low magnification overview. White rectangle highlights an OPC infected with ZIKV and a higher magnification view is shown in the right panel (arrow). Scale bar 20µm. **d-e**, Primary human astrocytes are vulnerable to ZIKV infection. **d**, overview of SP region infected with ZIKV. Arrows indicate infected astrocytes. Scale bar 20 µm. **e**, High magnification example of a ZIKV infected astrocyte. Scale bar 10 µm. **f**, Images show representative pattern of ZIKV infection of hPSC-derived astrocytes (see Methods). Scale bar 100 µm. Bar chart shows percentage of ZIKV-BR and ZIKV-PR infected cells cultured in the presence of IgG control or anti-AXL blocking antibody, relative to no treatment, showing a 96% or 70% decrease for ZIKV-BR and ZIKV-PR respectively (*n* = 4 for each strain, *p* = 0.0003 or *p* = 0.0016 by 2-way ANOVA for ZIKV-BR and ZIKV-PR respectively).

Based on the high infectivity of astrocytes in the cortical plate and the clinical findings of cortical plate abnormalities, we further analyzed mechanisms of viral entry in human astrocytes. Recent experiments demonstrated that blocking AXL in skin and lung epithelial cell culture substantially reduced ZIKV entry^11^ and blocking AXL in astrocytes similarly reduced dengue virus entry^16^. To test the hypothesis that AXL receptor is involved in mediating ZIKV entry into human astrocytes, we cultured astrocytes derived in vitro from human pluripotent stem cells (hPSCs)^17,18^ and infected them with ZIKV. At 2 hpi, we observed increased protein levels of AXL and the autophagy marker LC3B in astrocyte lysates as with lysates of placental CTBs at 72 hpi (**Extended Data** Fig. 7). We next pre-incubated the astrocytes with an antibody that blocks the extracellular domain of AXL before infecting with ZIKV. We found that blocking AXL substantially reduced the infection of astrocytes at an MOI of 10 (Fig. 3f). The blocking antibody results are consistent with a model in which AXL contributes to virus entry into cells.

We next searched for pharmacological methods to prevent or diminish the effects of ZIKV infection in human cells through a cytopathic effect (CPE)-based assay. We used U87 cells, a glioblastoma line that expresses high levels of astrocyte and radial glia marker genes^19^. U87 cells infected with ZIKV show a profound CPE at 48 hpi, with significant cell death occurring at 72 hpi (data not shown). Recent reports suggested that chloroquine, a drug historically used to treat or prevent malaria, could protect cells against ZIKV-induced death^20^. We observed a modest protective effect against ZIKV-induced CPE using chloroquine at 2-10 µM, but observed dramatic toxicity of the drug at over 10 µM (data not shown).

We further examined compounds from the macrolide class for protective effects. This class of drugs includes widely used antibiotics that are safe during pregnancy^21^, and certain macrolide antibiotics have been documented to block flavivirus replication^22^. By monitoring viral envelope production, we found that a commonly used macrolide, azithromycin (AZ), dramatically reduced ZIKV-induced CPE in U87 cells (Fig. 4a), and inhibited viral proliferation thereby limiting infection in U87 cells and in hPSC-derived astrocytes (Fig. 4b and c, **Extended Data** Fig. 8). These effects were observed at an EC50 of 6 µM for cell viability, or 3-8 µM for inhibition of viral proliferation. These concentrations are comparable to those measured in placental tissues following maternal dosing of AZ^23,24^and in adult brain tissue^25^. Although the mechanism of action remains unknown, the inhibition of ZIKV by AZ may serve as a launching point toward finding additional potent and safe therapeutics in the context of pregnancy.

**Figure 4.**
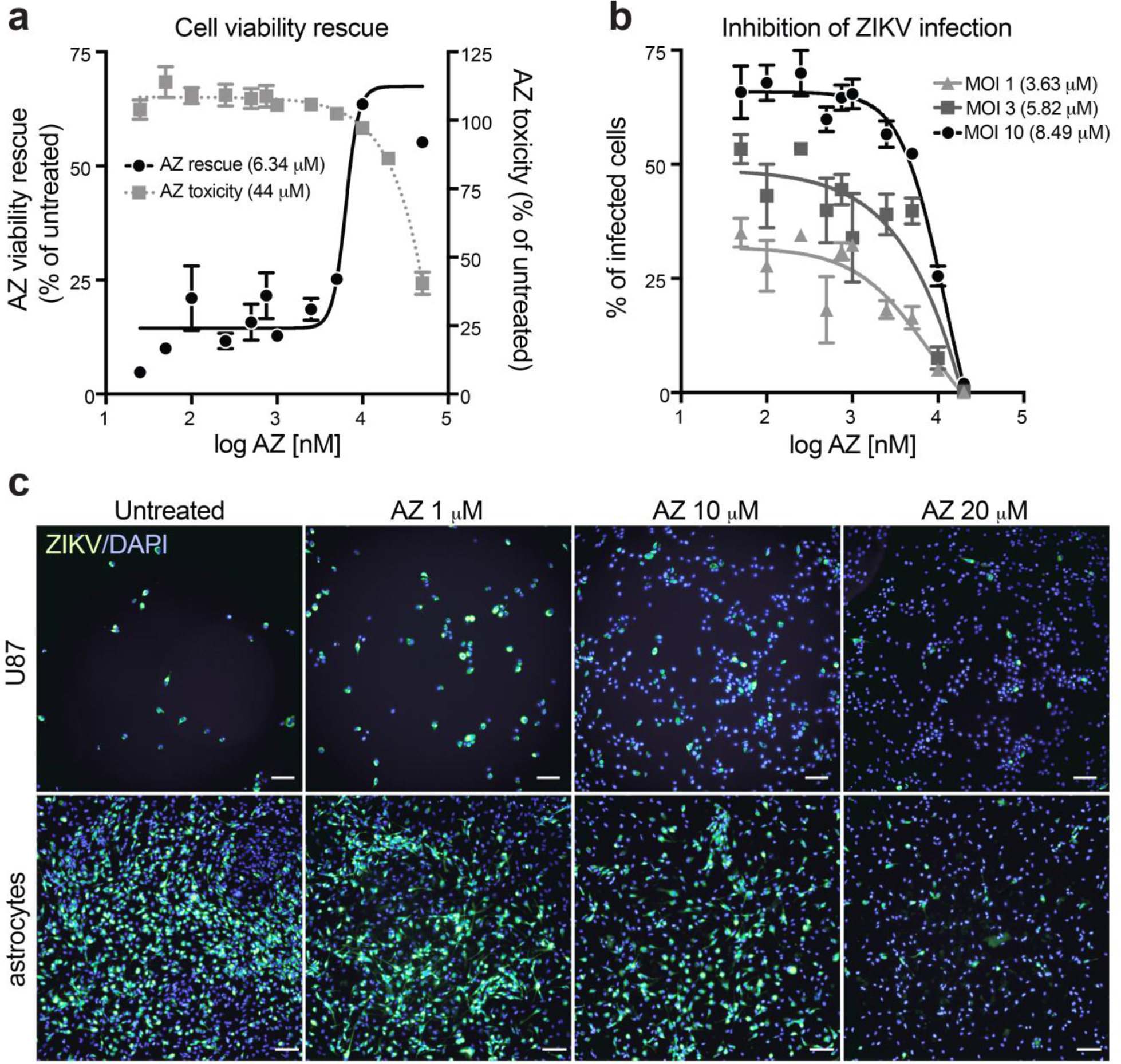
Azithromycin treatment prevents the ZIKV-induced cytopathic effect. **a**, Rescue of cell viability with azithromycin (AZ). U87 cells were pre-treated with AZ for 1 h and then infected with ZIKV-PR in the presence of AZ at an MOI of 10 for 72 h. AZ toxicity was evaluated in parallel in uninfected U87 cells treated with increasing concentrations of the drug. EC50 values for AZ-mediated cell viability rescue or AZ-mediated toxicity were 6.34 µM (*n* = 2) or 44 µM (*n* = 3), respectively. Error bars: SEM. **b**, AZ inhibits ZIKV infection/proliferation. U87 cells were treated with increasing concentrations of AZ and infected with increasing MOIs of ZIKV. The percentage of infected cells was calculated after immunostaining for anti-flavivirus envelope protein at 48 hpi. Half-maximal responses to AZ treatment were seen at 3.63 µM, 5.82 µM, or 8.49 µM for MOIs of 1, 3 or 10, respectively (*n* = 3 per MOI). Error bars: SEM. **c**, U87 cells (upper panels) and hPSC-derived astrocytes (lower panels) were treated with AZ and infected as indicated in **b**. At 48 hpi cells were immunostained for anti-flavivirus envelope protein and cellular DNA (DAPI). Scale bar 100 µm.

Together, our work highlights cell type specific patterns of ZIKV infection in the first and second trimester human placenta and developing brain, provides experimental evidence that AXL mediates ZIKV entry into relevant human brain cells, and suggests a candidate therapeutic agent that could be rapidly implemented to combat ZIKV pathophysiology in pregnant women. While preventative measures, such as mosquito abatement and a ZIKV vaccine, are imperative for long term control of this pathogen, small molecule therapeutics are also of great interest for prophylactic use or the treatment of acute infections in the context of pregnancy. We find that a common antibiotic, the macrolide azithromycin, has surprising activity against ZIKV through an unknown mechanism. Although the macrolides are known as antibacterial agents, previous work has shown that a related class of compounds, the avermectins, possess potent inhibitory activity against a range of flaviviruses, including yellow fever virus, dengue virus, and Japanese encephalitis virus^22^. Importantly, the macrolides are generally known to be safe in the context of pregnancy, which raises the possibility that oral macrolides could be used to mitigate or prevent the harmful effects of ZIKV infection in the unborn. Future work will be required to determine whether macrolides are capable of reducing ZIKV infection or replication in the critical tissues and cell types of the placenta and developing brain.

## Online Methods

### Cell lines

Vero and U87 cell lines were cultured in Dulbecco’s modified Eagle’s medium (DMEM) containing 10% fetal bovine serum, 2 mM L-glutamine, 100 U/ml penicillin-streptomycin, and 10 mM HEPES buffer at 37 °C with 5 % CO2. Human astrocytes (after 8 months of in vitro development) were derived from pluripotent stem cells (NIH Human Embryonic Stem Cell Registry line WA09 (H9) at passages 30-35) according to a recently published protocol^17^ and maintained in neural media composed of DMEM/F12 with sodium pyruvate and Glutamax, N2, B27, heparin and antibiotics. Media was either supplemented with growth factors (Epidermal Growth Factor (10 ng/ml) and Fibroblast Growth Factor (10 ng/ml)) or with Ciliary Neurotrophic Factor (10 ng/ml) during experiments. These cells show high levels of AXL expression (data not shown), in accordance with previously described transcriptional profiles of single fetal astrocytes^12^. hPSC-derived astrocytes were chosen due to their high fidelity to fetal astrocytes in vivo. All cell lines tested negative for mycoplasma using MycoAlert (Lonza).

### Virus propagation and titering

ZIKV, strains SPH2015 (Brazil 2015, ZIKV-BR), PRVABC59 (Puerto Rico 2015, ZIKV-PR), and FSS13025 (Cambodia 2010, ZIKV-CAM), were propagated in vero cells infected at an MOI of 0.01. Supernatants were collected at 72 and 96 hpi, clarified by centrifugation at 350 × g for 5 minutes, and filtered through a 0.45µm SFCA membrane. For mock infections supernatant was collected from uninfected vero cells and prepared by the same protocol used to make viral stocks. Virus was titered by plaque assay and focus assay. Briefly, plaque assays were performed using vero cells with a 0.7 % agarose overlay and processed five days post infection. Focus assays were performed on vero cells and processed 24 hpi with the anti-flavivirus group antibody (Millipore MAB10216, clone D1-4G2-4-15) at a dilution of 1:250. Titers determined by both methods were consistent. Each strain was sequence-verified using a previously published protocol^26^ and all viral stocks tested negative for mycoplasma contamination by MycoAlert (Lonza). ZIKV-PR and ZIKV-CAM continued to test negative after prolonged incubation in culture (96 h). Contamination of ZIKV-BR with mycoplasma was detected at low levels after 72-96 h in culture. No other evidence of contamination was seen in cells infected with this viral strain.

### Brain samples

De-identified primary tissue samples were collected with previous patient consent in strict observance of the legal and institutional ethical regulations. Protocols were approved by the Human Gamete, Embryo and Stem Cell Research Committee (institutional review board) at the University of California, San Francisco.

### Placenta samples

This study was approved by the UCSF Committee on Human Research. All donors provided written informed consent. The samples were processed immediately after they were acquired.

### Chorionic villus explants and isolated cytotrophoblasts

Chorionic villous explants were prepared and CTBs were isolated by using previously described methods^27,28^. Explants were cultured for 24 h prior to infection, which enabled attachment to a matrix substrate and extension of cell columns. Explants were incubated for 2 h with 5.6×10^5^ pfu ZIKV-BR, 5.6×10^6^ pfu ZIKV-CAM, or 1.1×10^6^ pfu ZIKV-PR. Purified CTBs were infected at an MOI of 1 within 1 h of cell plating and incubated with virus for 2 h. Following incubation with virus, the media was replaced with fresh media prior to incubation for 24 or 48 h. As a negative control, conditioned medium from uninfected vero cells was processed according to the protocol that was used to make viral stocks (mock infection). The number of samples that were incubated with each strain of virus and the results (as determined by immunostaining for anti-flavivirus group antigen, see below) are summarized in **Table 1**.

**Table 1.** Summary of results for *in vitro* placental infections. Data are shown as number of specimens cultured from different placentas/number in which infection was detected.

|  | ZIKV-CAM | ZIKV-PR | ZIKV-BR |
| --- | --- | --- | --- |
| First trimester explants | 5/5 positive | 3/3 positive | 3/3 positive |
| Second trimester explants | 3/3<br>(positive, but weaker staining and fewer foci) | 3/3<br>(positive, but weaker staining and fewer foci) | 1/1<br>(positive, but weaker staining and fewer foci) |
| First trimester isolated cytotrophoblasts | 3/3 positive | 2/2 positive | 2/2 positive |
| Second trimester isolated cytotrophoblasts | 5/5 positive | 3/3 positive | 5/5 positive |

### Developing brain organotypic slice culture experiments

Human primary cortical tissue blocks were embedded in 3.5% low melting point agarose (Thermo Fisher) and sectioned perpendicular to the ventricle to 300 µm using a Leica VT1200S vibrating blade microtome in artificial cerebrospinal fluid containing 125 mM NaCl, 2.5 mM KCl, 1 mM MgCl_2_, 1 mM CaCl_2_, 1.25 mM NaH_2_PO4. Slices were transferred onto slice culture inserts (Millicell) in 6- well culture plates (Corning) and cultured in media containing 66% Eagle’s Basal Medium, 25% Hanks Balanced Salt Solution, 5% Fetal Bovine Serum, 1% N-2 supplement, 1% penicillin/streptomycin, and glutamine (Thermo Fisher). Slices were cultured in a 37 °C incubator at 5% CO_2_, 8% O_2_ overnight. Slices were then incubated with 2.2×10^6^ pfu ZIKV-BR, 1.1×10^7^ pfu ZIKV-CAM, or 2.2×10^7^ pfu ZIKV-PR for 4 h, before replacement with fresh media and culture for an additional 72 hpi. Tissue samples were fixed overnight in 4% paraformaldehyde.

### Placental tissue immunohistochemistry

Placental tissue and explants were fixed, embedded and sectioned as described previously^29^. CTBs, attached to the tissue culture dish or transwell filters were fixed for 10 minutes in 3.7% paraformaldehyde and permeabilized with cold methanol. Double immunolocalization was performed as described previously^29^. Primary antibodies (mouse mAbs unless stated otherwise) were used at the following dilutions: rabbit mAb anti-AXL (10 ng/ml, Cell Signaling 8661), anti-CD68 (1:100, Dako, Clone KP1), anti-CK (1:200, 7D3 rat mAb, made in our laboratory^30^), anti-flavivirus group antigen antibody (1:100, EMD Millipore MAB10216), or rabbit pAb anti-vWF (1:100, Novus Biologicals NB600-586). Binding of the primary antibodies was detected via FITC-conjugated secondary antibodies (anti-mouse or anti-rabbit) or rhodamine-conjugated anti-rat (Jackson ImmunoResearch) diluted 1:100. The samples were imaged by confocal microscopy (Leica SP5) or by using an epifluorescence microscope (Leica DM 5000B with a 350 FX camera).

### Brain tissue immunohistochemistry

Heat-induced antigen retrieval was performed in 10 mM sodium citrate buffer, pH 6 at 95 °C for 20 minutes. Slices were incubated in blocking buffer containing 10% Donkey Serum, 1% TritonTM-X100 and 0.2% gelatin diluted in PBS at pH 7.4 for 1 hour. Primary antibodies: mouse anti-flavivirus group antigen (1:100, EMD Millipore MAB10216), goat anti-SOX2 (1:250, Santa Cruz SC17320), rabbit anti-SATB2 (1:200, Abcam SC81376), chicken anti-GFAP (1:500, Abcam ab4674), rabbit anti-IBA1 (1:200, Wako 019-19741), rabbit anti-OLIG2 (1:200, Millipore AB9610), rabbit anti-cleaved caspase 3 (1:100, Cell Signalling Techologies 9661), rabbit-anti PAX6 (1:200, Biolegend, previously Covance, PRB-278P), or rabbit-anti CD31 (1:200, Abcam ab28364) were diluted in blocking buffer as described above, and slices were incubated overnight at 4 °C. Binding was revealed using an appropriate Alexa FluorTM 488, Alexa FluorTM 546, Alexa FluorTM 594 and Alexa FluorTM 647 fluorophore-conjugated secondary antibody (Thermo Fisher) diluted 1:1000. Slices were incubated with secondary antibodies overnight at 4 °C, and cell nuclei were counter-stained using DAPI (Thermo Fisher). All washes were performed using PBS without calcium/magnesium containing 0.5% TritonTM ‑ X100. Slides were mounted with Fluoromount (Southern Biotech). Images were collected using a Leica TCS SP5 X or Leica TCS SP8 confocal microscope.

### Immunoblotting

Preparation of lysates (from chorionic villi, purified CTBs or cultured astrocytes) and immunoblotting was done according to methods that we published previously^29^. The blots were probed with: rabbit mAb anti-AXL (Cell Signaling 8661), 10 ng/ml for placental lysates, 1:1000 for astrocyte lysates; rabbit mAb anti-LC3B (Cell Signaling 3868), 1:100 for placental lysates, 1:1000 for astrocyte lysates; rabbit pAb anti-p62 (Cell Signaling 5114), 1:100; anti-β-actin (SIGMA A441) for astrocyte lysates, 1:50,000, or anti-β-actin (C4, Santa Cruz Biotechnology SC47778) for placental lysates, 1:100. Binding of the primary antibodies was detected via peroxidase-conjugated antibodies (anti-mouse or anti-rabbit; Jackson ImmunoResearch) diluted 1:5,000 to 1:20,000. Quantitative measurements of the immunoblot signals were made by using scanning densitometry and ImageJ software.

### Quantification of staining in human cortical sections

Images were quantified using Imaris v. 8.2 (Bitplane). For each biological replicate, one (13 pcw, 14 pcw, 15 pcw, 20 pcw) or two (22 pcw) representative slices were chosen for quantification. On each slice, one rectangle was drawn, spanning from VZ to pia and 1 mm in width. This rectangle was portioned into five counting boxes with height of either 1 mm or one fifth the distance from the ventricular surface to pia, whichever was smaller, and evenly distributed across the full height. With the location of each box defined this way, box 1 was directly adjacent to the ventricular surface, boxes 2 and 3 were typically found in the outer subventricular zone (OSVZ), box 4 typically covered the subplate (SP), and box 5 was located in the cortical plate (CP) directly adjacent to the pia. First, cells positive for anti-flavivirus envelope protein were identified based on cytoplasmic staining around a clear nucleus positive for DAPI. Then, envelope positive cells were scored for the nuclear expression of SOX2 or SATB2 (for 13 pcw, *n* = 259 cells; for 14 pcw, *n* = 347 cells; for 15 pcw, *n* = 172 cells; for 20 pcw, *n* = 192 cells; for 22 pcw, *n* = 271 and *n* = 179 cells for each slice). The percentage of envelope staining cells that were also positive for SOX2 or SATB2 was calculated by combining cells in all five boxes. In addition the percentage of envelope positive cells in each box relative to total number of envelope positive cells in the entire rectangle was displayed for each box and biological replicate.

### AXL blocking

Cells were treated with AXL blocking antibody (R&D Systems AF154) or goat IgG control (R&D Systems AB108C) at 100 µg/mL for 1 h prior to infection with ZIKV, washed once and cultured for 48 h before immunostaining against envelope protein and DAPI. Percentage of infected cells was calculated as described for azithromycin dose-response experiments (below).

### Cell viability assay

U87 cells were treated with specified concentrations of azithromycin (Sigma 75199), chloroquine (Sigma C6628), or with vehicle (PBS or DMSO final concentration <0.1%). One hour post treatment, cells were infected with ZIKV at an MOI of 1, 3, 10, or left uninfected. Cell viability was assayed using the CellTiter-Glo 2.0 luminescent assay (Promega) following the manufacturers recommendations. The percentage of viable cells in the presence of azithromycin was calculated as a fraction of uninfected/untreated cells (*n* = 3).

### Quantitative imaging for azithromycin dose response

U87 and hPSC-derived astrocyte cells were treated with specified concentrations of azithromycin for 1 h, then infected with ZIKV-PR or ZIKV-CAM at an MOI of 1, 3, or 10 (*n* = 2). Cells were fixed with 3.7 % PFA at 48 hpi, and stained with primary antibody mouse anti‑flavivirus envelope (1:250, EMD Millipore, MAB10216) and secondary goat anti‑mouse ALEXA FluorTM 488 (U87s) (1:2000, Thermo Fisher Scientific A‑11029) or secondary goat anti-mouse ALEXA‑FluorTM 594 (astrocytes) (1:2000 Thermo Fisher Scientific, A‑11032), and DAPI (Thermo Fisher). Two fields of view for each sample were imaged at 10X using a Nikon Eclipse Ti‑E microscope (Nikon Instruments) with an automated stage for high throughput imaging. The percentage of cells stained for ZIKV envelope protein was calculated using automated cell counting in Imaris v.8.2 (Bitplane).

### Evaluation of virus infectivity and infectious virus production

U87 cells were pre-treated with azithromycin (5, 10 or 20 µM) or vehicle for 1 h and infected with ZIKV-PR at an MOI of 3. The conditioned media of infected cells was collected at 1h, following removal of the inoculum, and at 24 or 48 hpi. The supernatant was clarified by filtering (0.45 µm PVDF filter), diluted in complete media and used to infect naïve cells. The percentage of infected cells after challenge with the supernatant from cells infected with ZIKV in the presence/absence of azithromycin was determined by viral envelope staining as described above.

To determine the rate of cell infection at 48 hpi in presence of azithromycin, U87 cells infected with ZIKV-PR at an MOI of 3 were dissociated with trypsin, fixed with 3.7% formaldehyde and permeabilized with 90% cold methanol overnight. Cells were then stained with mouse anti‑flavivirus envelope primary antibody (1:100 EMD Millipore, MAB10216), and secondary goat anti‑mouse ALEXA Fluor-TM 488 (1:1000 Thermo Fisher Scientific, A‑10680). The percentage of cells stained for ZIKV envelope protein was calculated by flow cytometry.

## Acknowledgements

This work was supported by the NIH/NINDS grant R01NS075998 and U01 MH105989 as well as a gift from Bernard Osher (to A.R.K.), NIH/NICHD grants P50HD055764 and R37HD076253 (to S.J.F), Howard Hughes Medical Institute (to J.L.D), NIMH (R01MH099595-01) and Paul G. Allen Family Foundation Distinguished Investigator Award (to E.M.U.), and Damon Runyon Cancer Research Foundation postdoctoral fellowship to A.A.P. (DRG-2166-13). The authors wish to thank Marc and Lynne Benioff for their financial support for these studies. We thank Robert Tesh (UTMB), Nikos Vasilakis (UTMB), Julio Rodriguez-Andres (CSIRO), Graham Simmons (BSRI), Charles Chiu (UCSF), Dan Lim (UCSF), John Liu (UCSF), Shaohui Wang (UCSF), Diego Acosta-Alvear (UCSF) and the Small Molecule Discovery Center for providing reagents and advice.

## Contributions

S.J.F. and H.R. conceived of and designed the experiments that employed placental cells. Y.Z. prepared the chorionic villus explants. Y.Z. and K.O. performed immunostaining on the placental samples, which Y.Z., K.O., and M.G. imaged. K.O. and Y.Z. performed the villus and cytotrophoblast immunoblotting experiments. S.J.F., Y.Z., K.O., and M.G. analyzed data from the placenta tissue experiments. H.R. and E.D.L. conceived of and designed the brain slice culture experiments. E.D.L., C.S.E., and W.R.M.L. performed the slice culture experiments as well as immunostaining, imaging, and image analysis. H.R. performed quantitative analysis of brain infections. E.D.L. conceived of the AXL in vitro blocking experiments which H.R. performed. J.S performed imaging, image analysis and immunoblots. T.J.N. and A.A.P. provided conceptual guidance throughout the project. K.A.K. made and titered viral stocks. H.R. performed infections on brain and placental tissues. C.A. and M.T.L. designed drug-response experiments with input from J.L.D., H.R. and K.A.K., which H.R., C.A., K.A.K., and M.T.L. performed and analyzed. R.K. derived astrocytes from hPSCs with guidance from E.M.U. Scientific guidance and supervision for the project was provided by S.J.F., J.L.D., and A.R.K. Manuscript was written by H.R., E.D.L., C.A., K.A.K., C.S.E., M.T.L., T.J.N., A.A.P., J.L.D., S.J.F, and A.R.K. with input from all authors.

## Extended Data

### Zika Virus in the Human Placenta and Developing Brain: Cell Tropism and Drug Inhibition

**Extended Data Figure 1.**
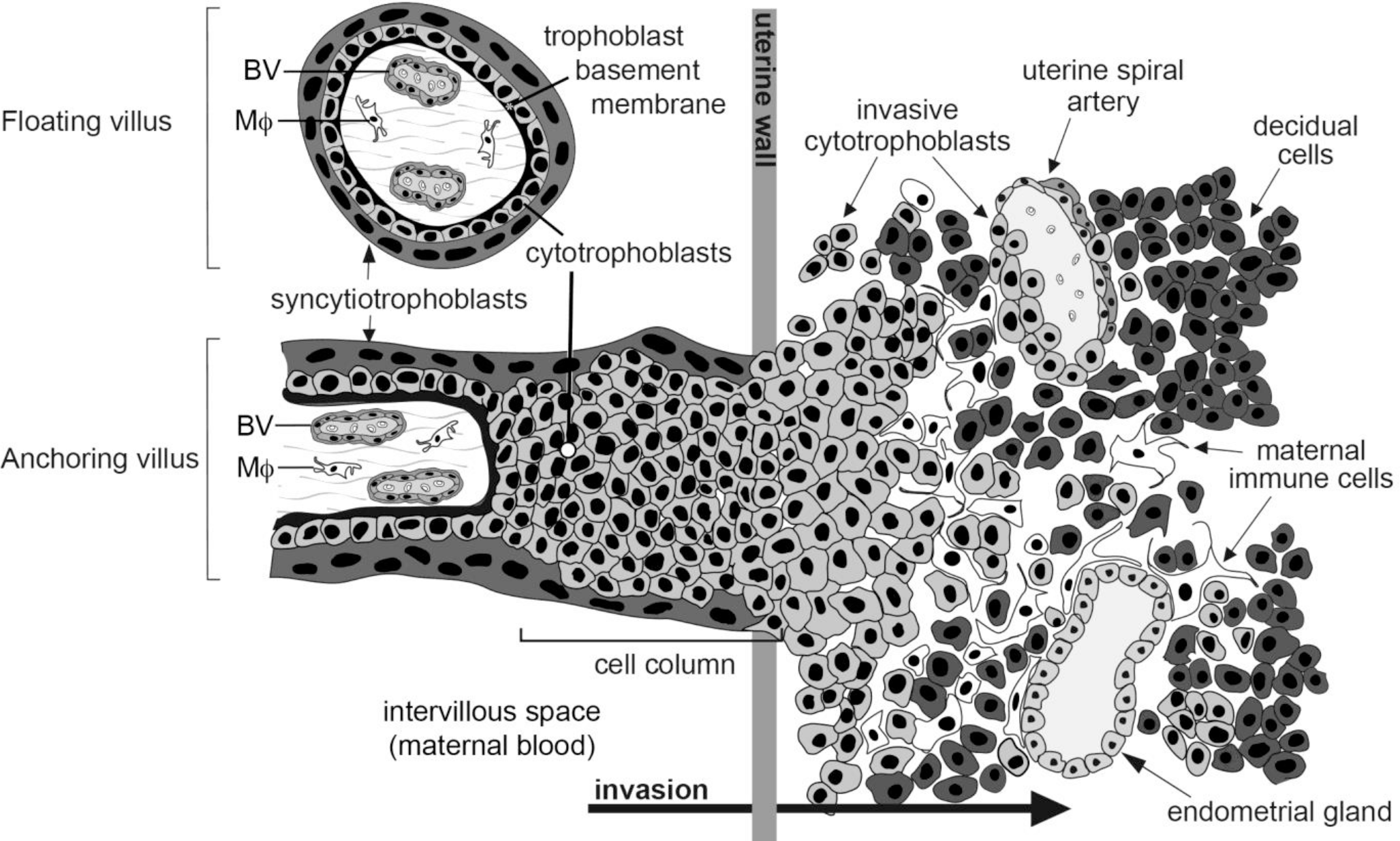
Sites where human placental trophoblasts interface with maternal cells (blood and uterine cells), and thus, where ZIKV infection might occur. Development of human placental chorionic villi entails the formation of two major trophoblast subtypes. Both differentiation processes involve cytotrophoblast (progenitors), a polarized layer of cells that is attached to the trophoblast basement membrane, which surrounds a stromal core rich in blood vessels (BV) that contains a specialized population of fetal macrophages (Mϕ; Hoffbauer cells). In one differentiation pathway, which gives rise to floating villi, cytotrophoblast progenitors fuse to form multi-nucleated syncytiotrophoblasts, which transport substances to and from maternal blood that perfuses the intervillous space in which they “float.” In the other differentiation pathway, which gives rise to anchoring villi, cytotrophoblasts form columns of nonpolarized cells that attach to and penetrate the uterine wall. The ends of the columns terminate within the superficial decidua, where they give rise to invasive (extravillous) cytotrophoblasts. During interstitial invasion, a subset of these cells, either individually or in small clusters, co-mingles with decidual, myometrial, and immune cells (primarily macrophages and NK cells). During endovascular invasion, masses of cytotrophoblasts breach and line the vessels, replacing the resident maternal endothelium and portions of the smooth muscle wall. These novel hybrid arteries, which are made up of maternal and embryonic/fetal cells, divert uterine blood flow to the placenta.

**Extended Data Figure 2.**
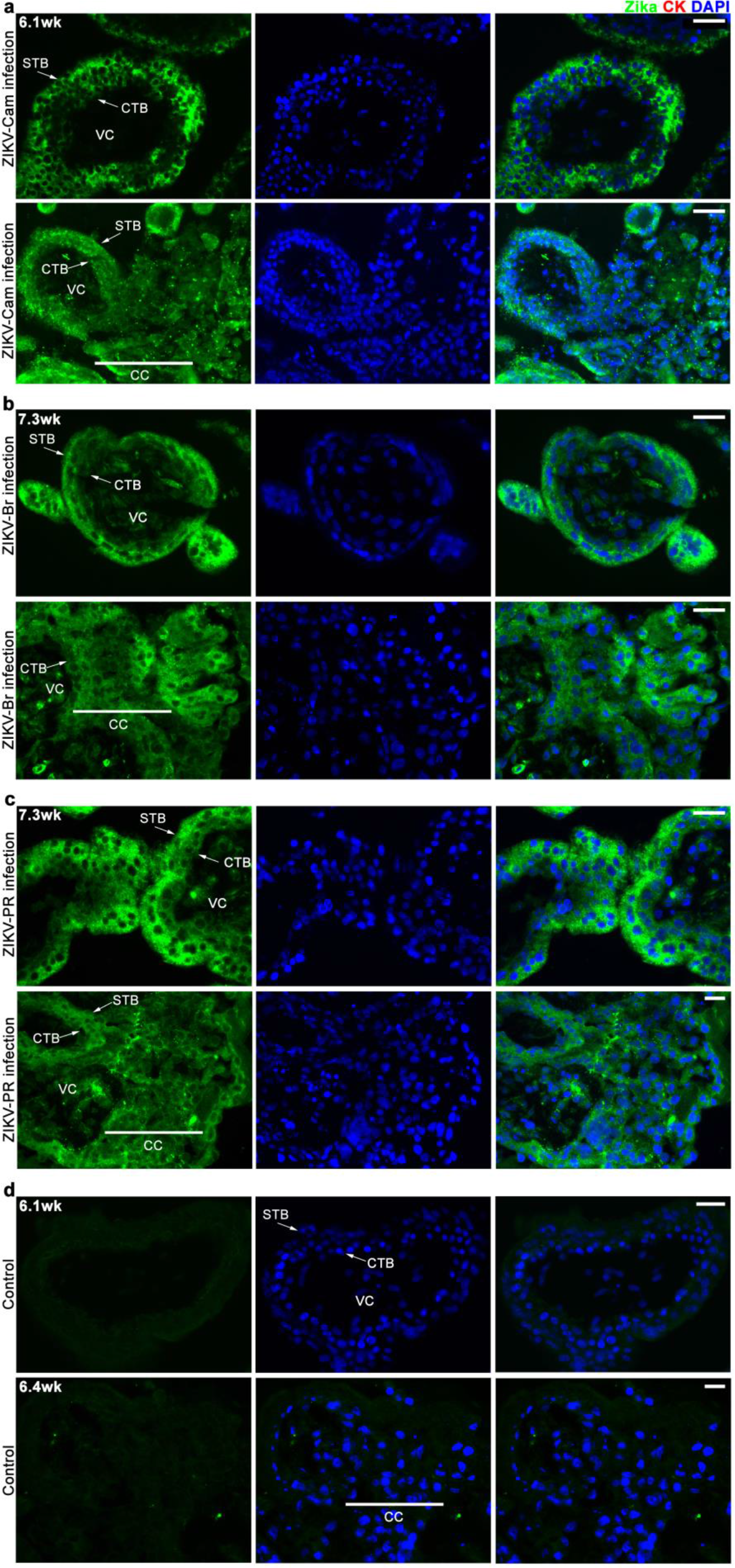
First trimester chorionic villus explants were susceptible to ZIKV-CAM, ZIKV-BR and ZIKV-PR infection. The explants were sectioned and stained with anti-flavivirus envelope protein (green). Nuclei were visualized with DAPI. For each panel, floating villi are shown in the top row and anchoring villi are shown in the bottom row. **a-c**, Syncytiotrophoblasts (STBs), cytotrophoblasts (CTBs), and cell column cytotrophoblasts (CCs) were susceptible to (**a**) ZIKV-CAM, (**b**) ZIKV-BR, and (**c**) ZIKV-PR infection. Images were acquired by fluorescence microscopy. **d**, Controls were mock infected. Scale bars, 25 µm.

**Extended Data Figure 3.**
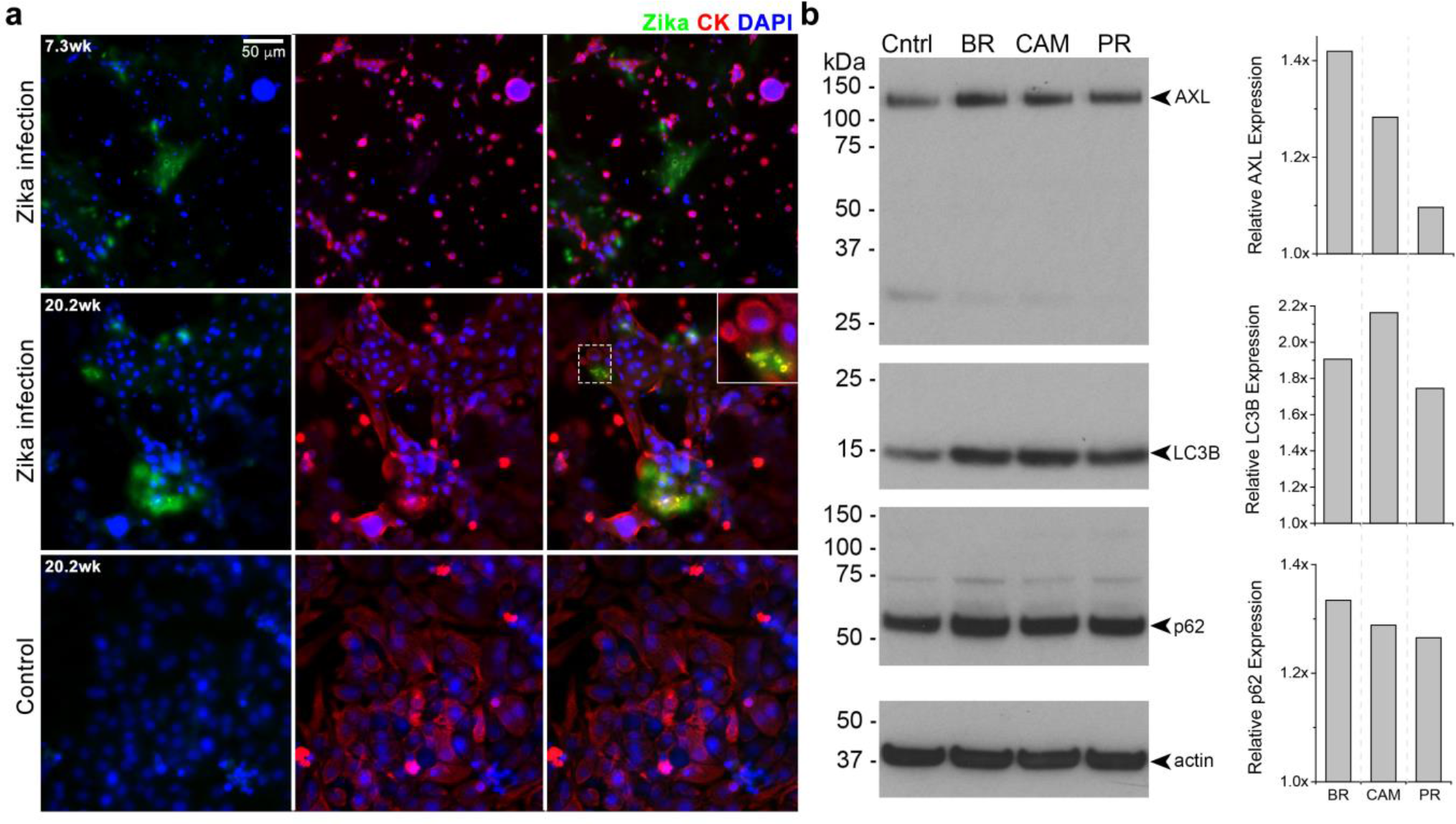
Cytotrophoblasts isolated from first and second trimester placentas were infected with ZIKV-CAM. Freshly isolated cells were plated and incubated with ZIKV within 1 h of plating. The cultures were stained with anti-flavivirus envelope protein (green) and anti-cytokeratin (CK; red) antibodies. **a**. At 24 hpi, infection of some cells was evident. As with the villus explants (Figure 1a), some areas of infectivity were associated with downregulation of CK staining (top row). Control cultures (bottom row) were mock infected. **b**. In cytotrophoblasts, infection was associated with up regulated expression of AXL, LC3B and p62. The quantification was relative to actin expression. Images were acquired by fluorescence microscopy. Inset = 2.5x

**Extended Data Figure 4.**
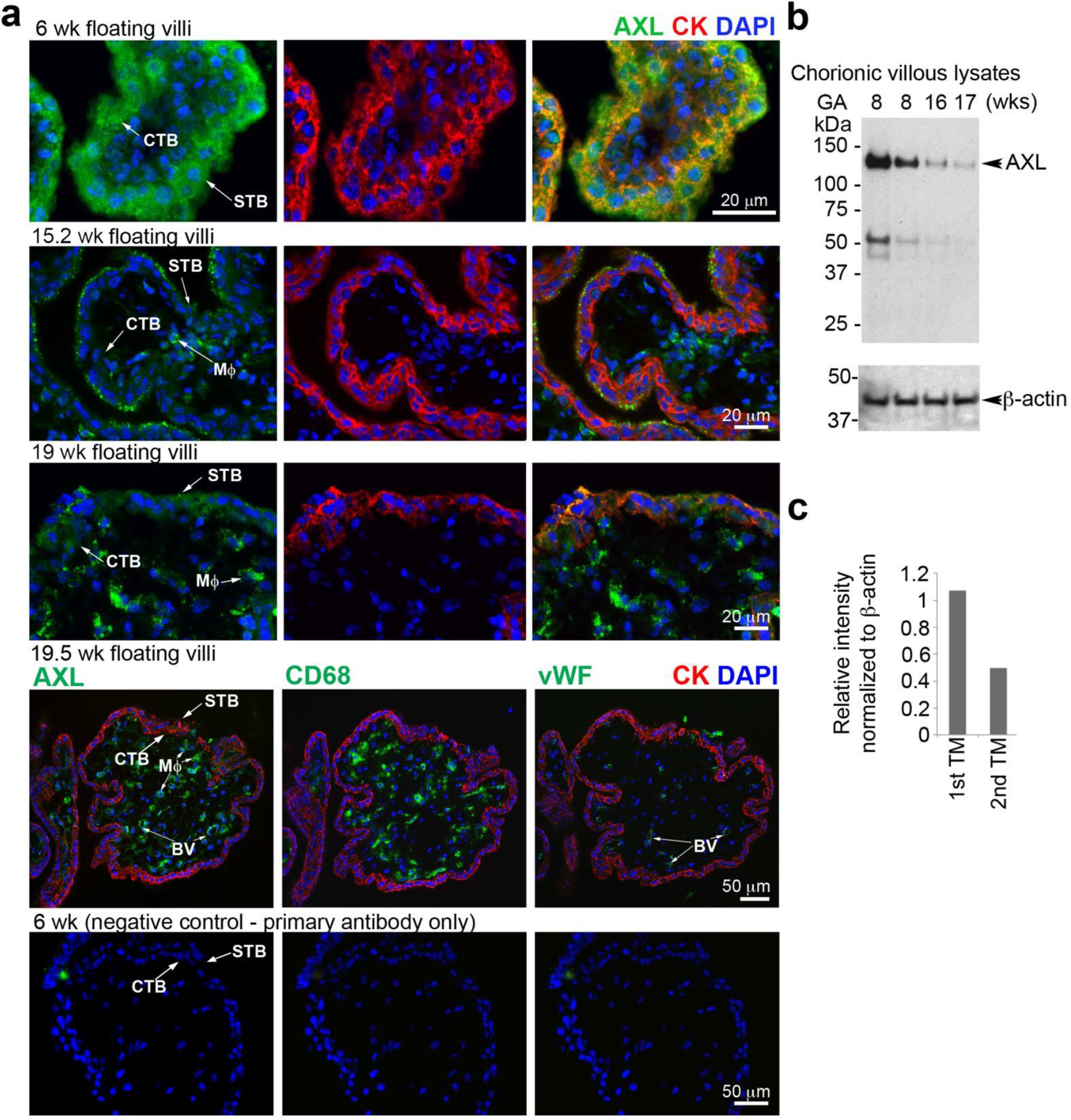
Immunolocalization of AXL in floating chorionic villi of the placenta revealed a strong signal during the first trimester, which was down regulated as gestation advanced. In all cases, staining with a rat anti-cytokeratin (CK; red) mAb (7D3; 7529679) confirmed trophoblast identity and DAPI was used to image nuclei. The pattern of AXL immunoreactivity (green) underwent striking changes as a function of gestational age. **a**, In first trimester floating villi (6 wk), an AXL signal was detected in association with STBs, including the syncytial surface that was in direct contact with maternal blood. The underlying layer of CTB progenitors also reacted with anti-AXL. By second trimester, TB immunoreactivity was greatly reduced. In some locations, a signal was detected in association with the STB surface (15.2 wk) and occasionally CTB progenitors stained weakly (19 wks). At this stage (19.5 wk), macrophages and blood vessels in the villous cores also reacted with anti-AXL. Their identity was confirmed via CD68 and von Willebrand Factor (vWF) expression, respectively. No signal was observed in tissue sections that were stained with anti-AXL alone (6 wk). **b,c**, Immunoblotting of lysates prepared from chorionic villi at different weeks of pregnancy confirmed this pattern of gestational age regulation, i.e., strongest expression in first trimester and down regulation during second trimester. The major band corresponded to the expected molecular weight of AXL (~144,000). An immunoreactive lower molecular weight band was also detected, perhaps a cleavage product generated in the proteinase-rich placental environment.

**Extended Data Figure 5.**
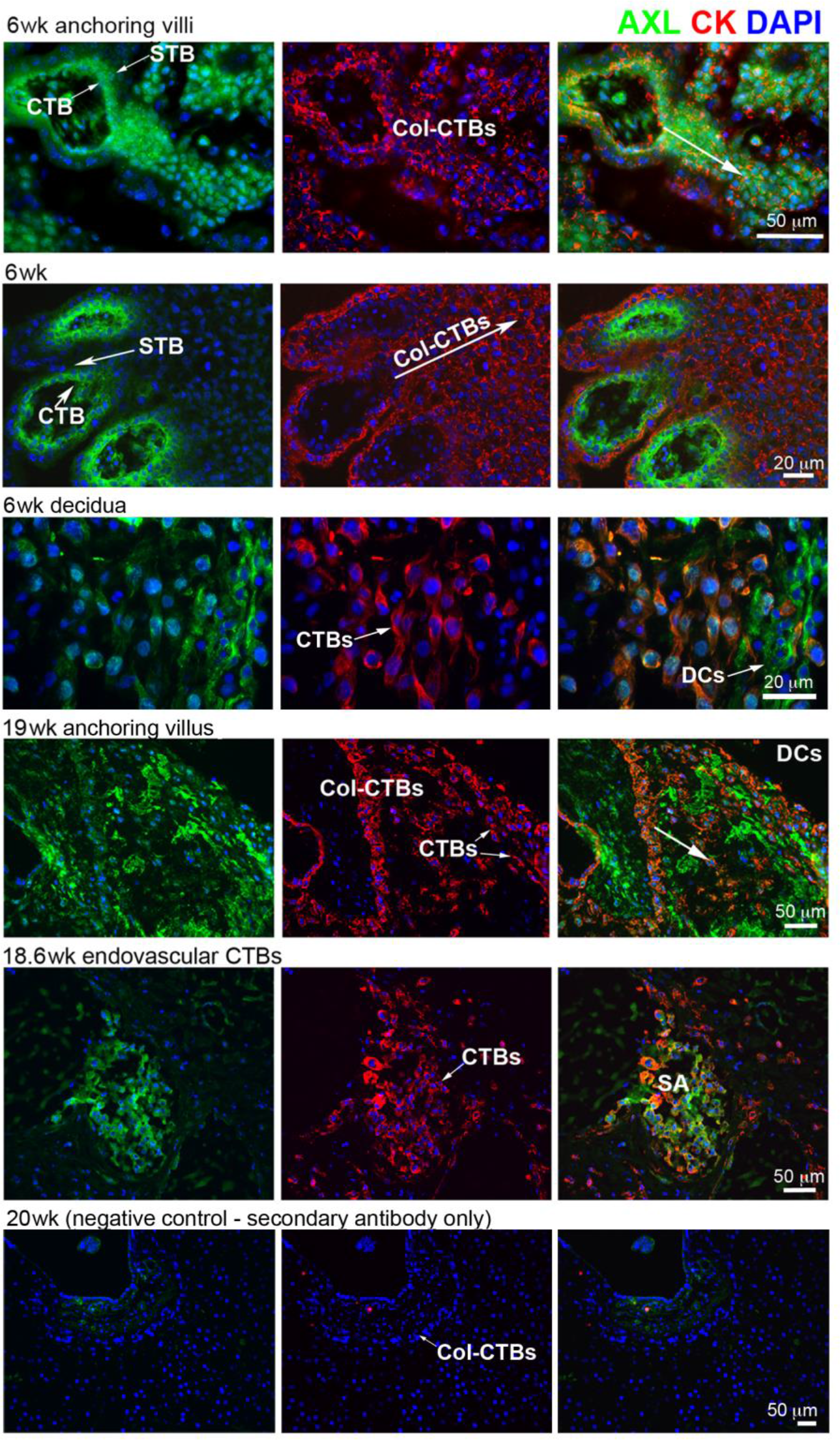
Immunolocalization of AXL in anchoring chorionic villi of the placenta revealed a strong signal during the first trimester, which was down regulated as gestation advanced. (First trimester; 6 wks) Two patterns were observed. In some anchoring villi, syncytiotrophoblasts (STB) and cytotrophoblasts (CTB) in the villi stained brightly as did the CTB cell columns (Col-CTB) that connect the placenta to the uterine wall. More deeply inside the uterus CTBs come into contact with decidual cells (DC), which also immunostained for AXL. (Second trimester; 19 wks, 18.6 wks). CTBs within the uterine wall had a less intense AXL signal and DCs continued to react with anti-AXL. During this period, CTB invasion and remodeling of maternal spiral arteries that traverse the uterus are robust. Immunostaining of these vessels showed AXL expression in association with endovascular CTBs that displaced maternal endothelium. No specific immunoreactivity was detected in the control conditions, staining with the primary (data not shown) or secondary antibody alone.

**Extended Data Figure 6.**
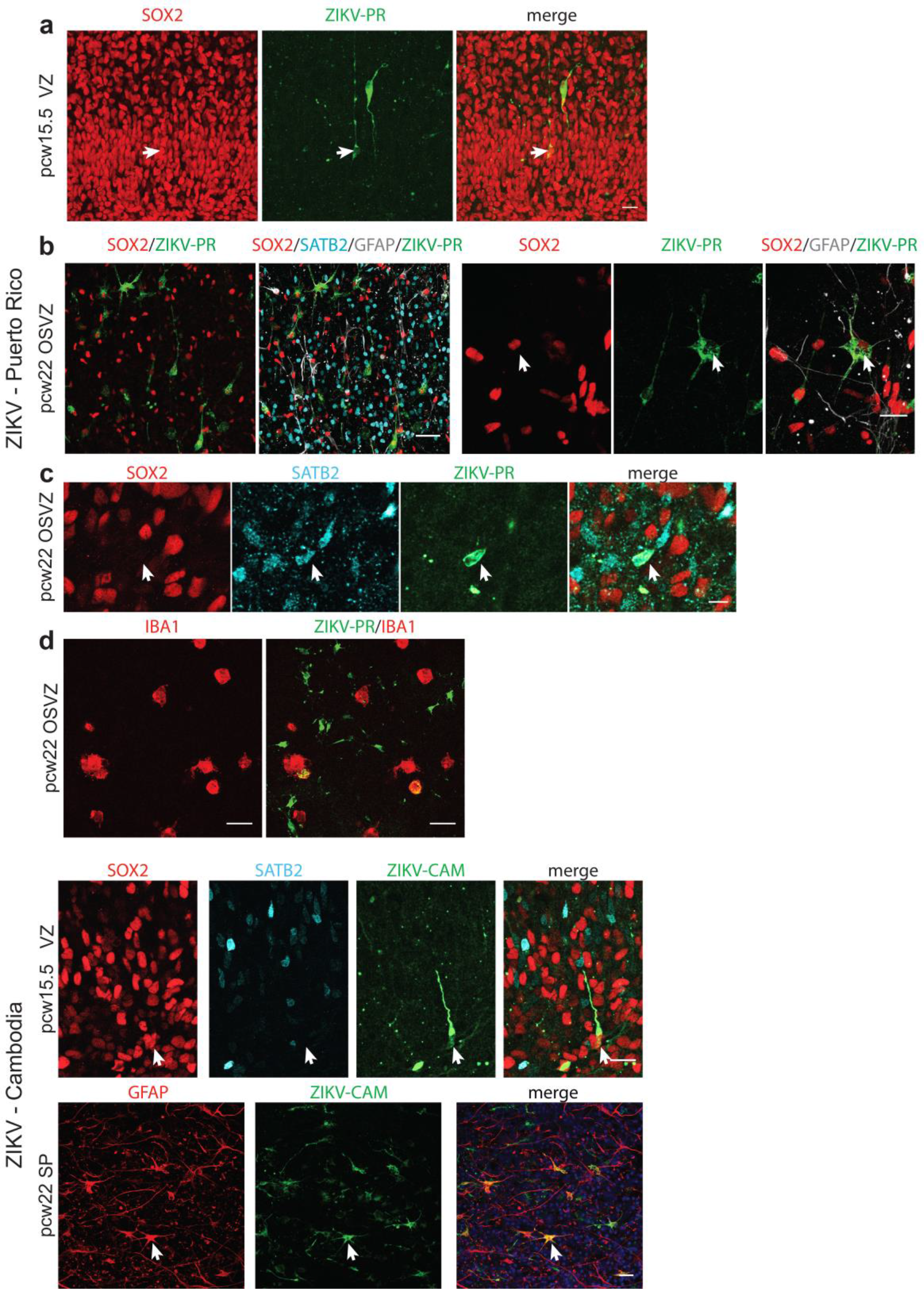
Cell tropism in the brain is conserved across ZIKV strains. **a-d**, Infection of human brain slices with ZIKV-PR followed by immunostaining for ZIKV envelope protein and cell type identity markers. ZIKV-PR infects radial glia (arrow) in the cortical VZ at mid-neurogenesis **(a)**, immature astrocytes (arrow) at late second trimester **(b)**, immature neurons (arrow, **c**), and microglia **(d)**, consistent with the patterns of infectivity of ZIKV-BR shown in Figure 2 and Fig. 3. **e-f**, Similar pattern of infectivity was observed for ZIKV-CAM. **e**, Images show example of a radial glia cell in the VZ at mid-neurogenesis infected with ZIKV-CAM. **f**, ZIKV-CAM infects immature astrocytes at mid-gestation (arrow). Scale bars: a, d, e, f ‑ 20µm, c ‑ 10µm b, 50µm

**Extended Data Figure 7.**
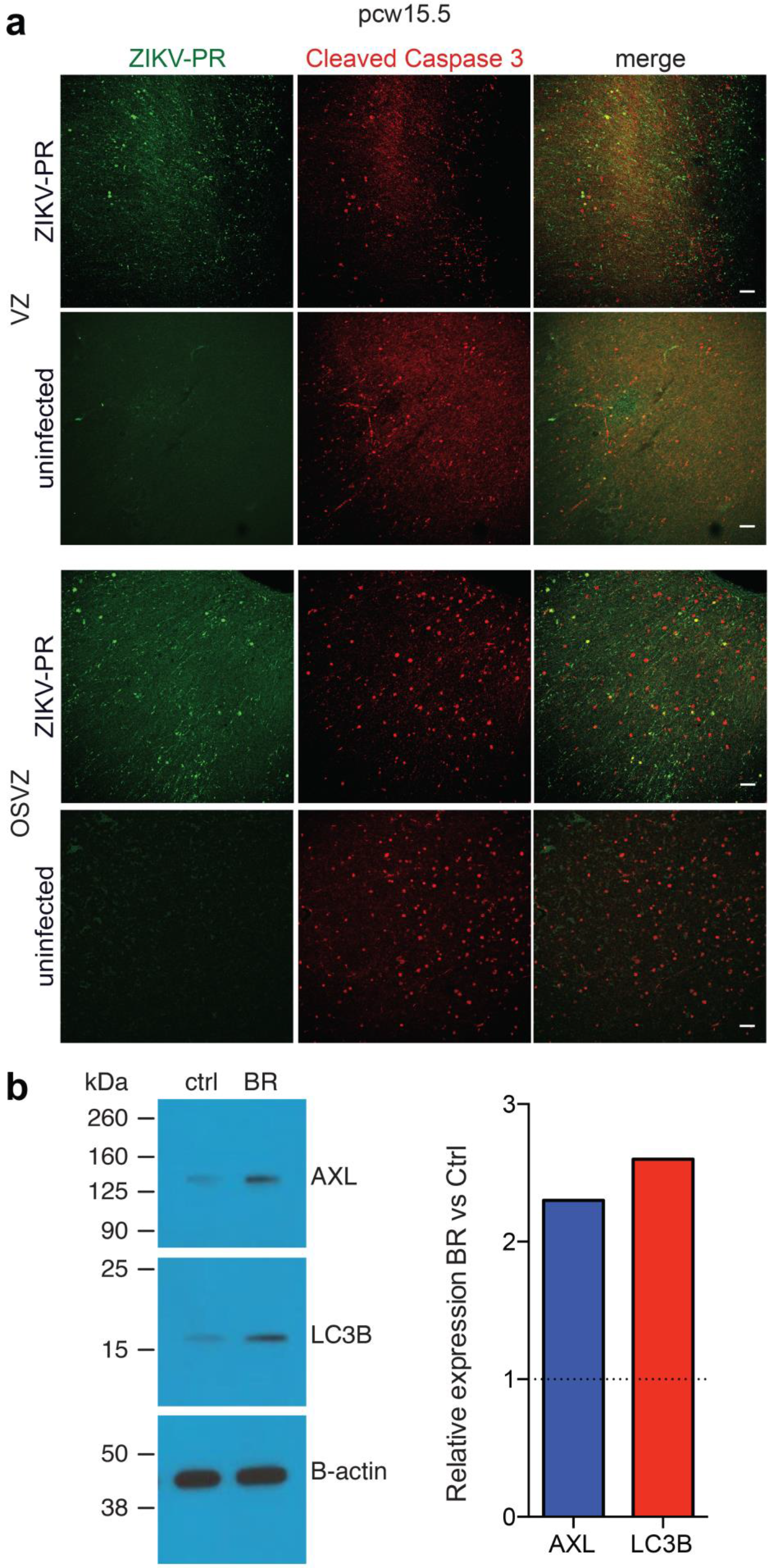
No increase in cell death in ZIKV-PR infected slices. **a**, Top 2 rows show cleaved caspase 3 staining in ZIKV-PR infected slice compared to mock-infected (“uninfected”) in the VZ. Bottom 2 rows show cleaved caspase 3 in the ZIKV-PR infected slides compared to mock-infected (“uninfected”) in the OSVZ. Merged image shows lack of overlap between ZIKV-PR infected cells and cleaved caspase 3+ cells. Scale bar 100µm. **b**, Increase in AXL protein and LC3B protein in astrocytes in vitro following ZIKV-BR infection. Immunoblotting of lysates prepared from hPSC-derived astrocytes 2 hours post infection with ZIKV-BR or MOCK (ctrl). Antibodies against AXL (144kDa) and LC3B (16kDa) were used to probe protein levels. Bar chart shows that following normalization with β-actin, a greater than 2-fold increase in AXL and LC3B protein level was detected.

**Extended Data Figure 8.**
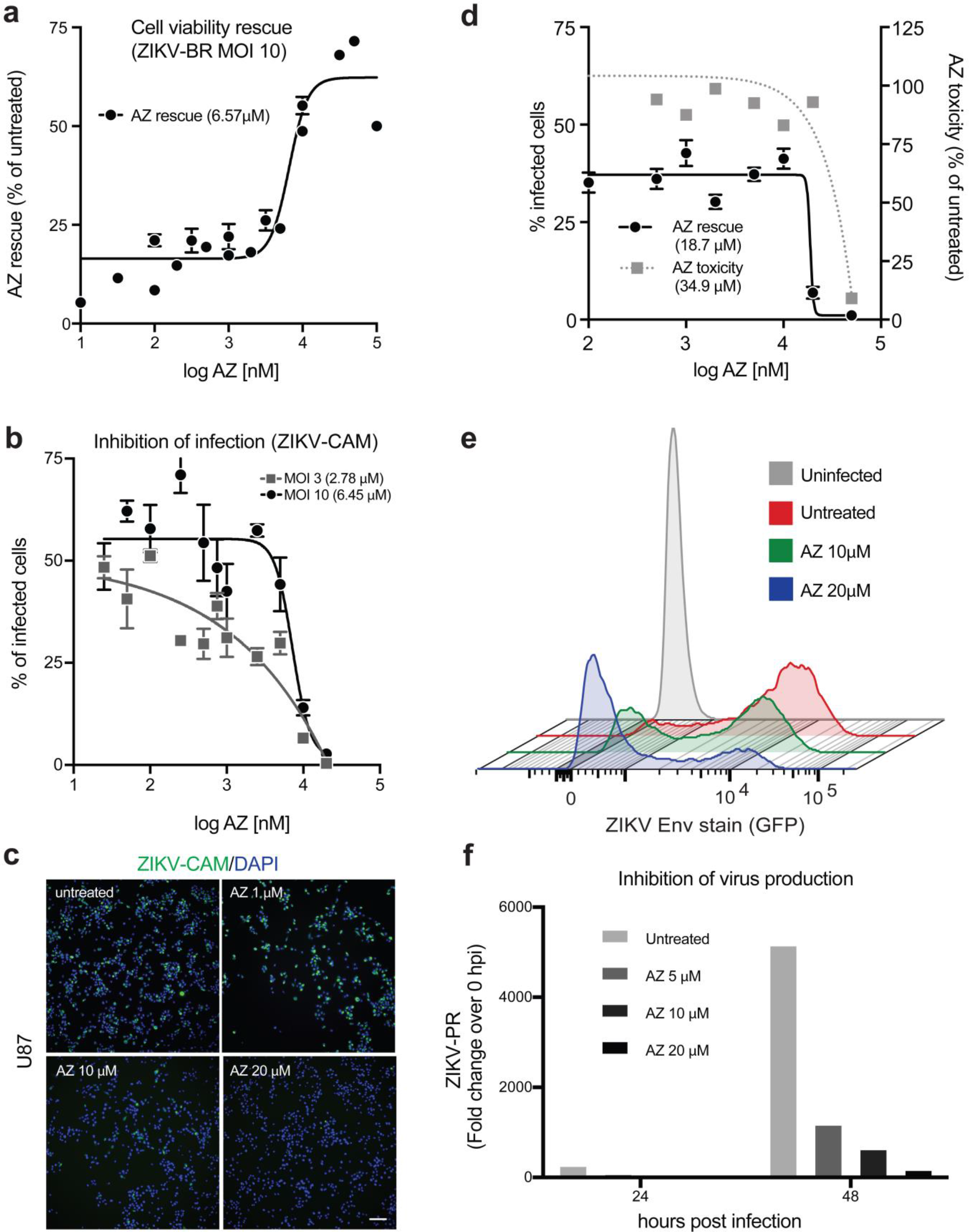
Azithromycin reduces the ZIKV-dependent cytopathic effect and inhibits viral proliferation of multiple viral strains. U87 cells (**a-c,e,f**) or hPSC-derived astrocytes (**d**) were pre-treated with azithromycin (AZ) for 1 h and then infected with the indicated strain of ZIKV at MOI 3 (**b,e**), or 10 (**a-d**) in the presence of AZ. **a**, AZ-mediated rescue of cell viability in cells infected with ZIKV-BR. The percentage of viable cells at 72 hpi with ZIKV-BR, in presence/absence of AZ, was determined using the CellTiter-Glo luminescent assay. EC50 values for AZ-mediated cell viability rescue with ZIKV-BR MOI 10 was 6.57 µM (*n*=3). Error bars: SEM. **b**, AZ inhibits infection/proliferation of ZIKV. U87 cells infected with ZIKV-CAM in presence/absence of AZ were immunostained for ZIKV envelope expression at 48 hpi. Half-maximal responses to AZ treatment were seen at 2.78 µM or 6.45 µM for MOIs of 3 or 10 respectively (*n*=3 per MOI). **c**, representative images from the infection/immunostaining described in b. (**d**), hPSC-derived astrocytes were infected with ZIKV-PR in presence of increasing concentrations of AZ. Drug-mediated toxicity of cells was evaluated in parallel. The percentage of infected cells was calculated following quantification of cells immunostained for ZIKV envelope (see Fig. 4). Half-maximal responses to AZ treatment or AZ-mediated toxicity (TC50) were 18.7 µM (*n*=4) or 34.9 µM (*n*=1) respectively. **e**, AZ prevents ZIKV infection/proliferation U87 cells treated with AZ and infected with ZIKV at an MOI of 3 were assayed for viral envelope protein immunostaining by flow cytometry (*n*=10000 cells/condition). **f**, AZ treatment reduces infectious ZIKV particle production. Virus production from U87 cells infected with ZIKV-PR at an MOI of 3 in presence/absence of AZ, at 24 h and 48 h post-infection, was determined by titering onto naive cells. Percentage of infection in the target cells was determined by ZIKV envelope staining (*n*=1 per AZ concentration).

